# Targeting the SHP2 phosphatase promotes vascular damage and inhibition of tumor growth

**DOI:** 10.1101/2020.10.06.327882

**Authors:** Yuyi Wang, Ombretta Salvucci, Hidetaka Ohnuki, Andy D. Tran, Taekyu Ha, Jing-Xin Feng, Michael DiPrima, Hyeongil Kwak, Dunrui Wang, Michael Kruhlak, Giovanna Tosato

## Abstract

The tyrosine phosphatase SHP2 is oncogenic in cancers driven by receptor-tyrosine-kinases, and SHP2 inhibition reduces tumor growth. Here, we report that SHP2 is an essential promoter of endothelial cell survival and growth in the remodeling tumor vasculature. Using genetic and chemical approaches to inhibit SHP2 activity in endothelial cells, we show that SHP2 inhibits pro-apoptotic STAT3 and stimulates proliferative ERK1/2 signaling. Systemic SHP2 inhibition in mice bearing tumors selected for SHP2-independent tumor-cell growth, promotes degeneration of the tumor vasculature and blood extravasation; reduces tumor vascularity and blood perfusion; and increases tumor hypoxia and necrosis. Reduction of tumor growth ensues, independent of SHP2 targeting in the tumor cells, blocking immune checkpoints or recruiting anti-tumor macrophages. We also show that inhibiting the Angiopoietin/TIE2/AKT cascade magnifies the vascular and anti-tumor effects of SHP2 inhibition by blocking tumor endothelial AKT signaling, not a target of SHP2. Since the SHP2 and Ang2/TIE2 pathways are active in vascular endothelial cells of human melanoma and colon carcinoma, SHP2 inhibitors alone or with Ang2/Tie2 inhibitors hold promise to effectively target the tumor endothelium.

## Introduction

Endothelial cells, which line the internal surface of blood vessels, are structural components of the vasculature that supplies blood to tissues and are poised to respond to environmental insults. Considerable work has focused on targeting the structural functions of endothelial cells, particularly to limit the supply of blood and nutrients to cancer tissues. Neutralization of vascular endothelial growth factor (VEGF) reduces tumor neovascularization and drugs that neutralize VEGF activity have increased the survival for patients suffering from certain cancer types (1). However, resistance to anti-VEGF treatment is common and its emergence has been attributed to the presence of endothelial cell populations which are VEGF receptor-deficient, display ligand-independent VEGF receptor activity, rely on growth factors other than VEGF and to other causes (1-3). One such factor is Angiopoietin-2 (Ang2), and the combined neutralization of VEGF and Ang-2 has shown promising results in initial trials (1,4).

The structurally and functionally disordered tumor vasculature undergoes sustained remodeling in response to changing signals from cancer cells and the tumor microenvironment (5). In mice, anti-VEGF responsive “sprouting” tumor vessels evolve into anti-VEGF unresponsive arterial-venous “vascular malformations” (6). However, little is known about the mechanisms that orchestrate remodeling of post-angiogenic tumor vessels and how remodeling relates to anti-angiogenic therapy resistance.

EphrinB2 is a transmembrane ligand for Eph receptors that plays pivotal roles in vascular biology during development and after birth (7,8). Once tyrosine phosphorylated by either EphB receptor engagement or trans-phosphorylated by tyrosine kinase (TK) receptors, such as TIE2, FGFR and PDGFR, EphrinB2 regulates cell-to-cell adhesion and cell movement (8,9). We have identified a pro-survival role for tyrosine phosphorylated EphrinB2, which relies upon the associated phosphatase SHP2 (SRC homology 2-containing Protein Tyrosine Phosphatases (PTP)) preventing activation of a STAT-dependent pro-apoptotic pathway in endothelial cells (10). Inactivation of this pro-survival pathway is critical for the physiologic involution of hyaloid vessels in the developing eye (10).

We now hypothesized that EphrinB2/SHP2-dependent signaling plays a role in the regulation of tumor vessel survival. Here, we identify active SHP2 as an essential guardian of endothelial cells and tumor vessel persistence by simultaneously repressing STAT3 signaling and activating ERK1/2 signaling in endothelial cells. By rigorously selecting or genetically engineering tumors composed of SHP2 growth-independent tumor cells, we find that specific SHP2 inactivation promotes endothelial cell death and the involution of tumor vessels resulting in reduced tumor growth, without impacting the resting vasculature of normal tissues. The current identification of SHP2 as a previously unrecognized regulator of the tumor vasculature delineates a novel approach to reducing tumor vascularity and tumor growth.

## Materials and Methods

### Cells, cell culture and materials

HUVEC (Lifeline Cell Technology, Frederick MD; FC-0003), HDMEC (Clonetics, CC-2543), BMEC (gift from Dr. Jason M. Butler, Weill Cornell Medical College, New York, USA); the murine melanoma B16F10 (ATCC CRL-6475), Lewis lung carcinoma LLC1 (ATCC CRL-1642), mammary cancer 4T1 cell line (ATCC CRL-2539), plasmacytoma MOPC315 (ATCC TIB-23), colon adenocarcinoma MC-38 (a gift of Dr. James Hodge); and human lung carcinoma A549 (ATCC CCL-185), mammary gland carcinoma Hs 578T (ATCC HBT-126), osteosarcoma G-292 (ATCC CRL-1423), NUGC3 gastric adenocarcinoma (JCRB 0822), colon carcinoma RKO (ATCC CRL-2577) and HT29 (ATCC HTB-38), and embryonic kidney 293T (ATCC CRL-3216) were propagated as described in Supplementary Materials and Methods. All cells lines tested *Mycoplasma*-negative but were not authenticated in the laboratory. The inhibitors: SHP099 (Investigational Drugs Repository, DCTD, NCI and MedChemExpress, HY-100388), AMG386 (Amgen, Inc. under a Materials Cooperative Research and Development Agreement), Tofacitinib (Millipore Sigma, PZ0017), Ruxolitinib (Millipore Sigma, ADV390218177), PD098059 (MedKoo Biosciences, 401680) and SCH772984 (Selleckchem S7101) were used as detailed in Supplementary Materials and Methods.

### Gene silencing and expression

We generated lentiviral shRNA particles for silencing mouse EphB4 (MISSION shRNA; Sigma-Aldrich TRCN0000023619 and TRCN0000023621); SHP2 (MISSION shRNA; Millipore-Sigma, TRCN0000327987, TRCN0000029877, TRCN0000328059, TRCN0000029875, TRCN0000029878 and TRCN0000327984); controls pLKO (SHC001, no insert) and non-mammalian shRNA (Sigma; SCH002) in 293T cells using the third-generation lentiviral packaging system (10,11). Infected cells were selected with puromycin (1μg/ml, ThermoFisher, A11138) for 10 days. RNA purification, cDNA production and real-time PCR are described in Supplementary Materials and Methods.

### Flow cytometry and cell proliferation

Cell viability and cell cycle were measured by flow cytometry as described in Supplementary Materials and Methods. Cell proliferation was measured by ^3^H-thymidine incorporation as detailed in Supplemental Materials and Methods.

### ELISAs and cell death proteomic assay

Ang2 was measured by Mouse/Rat Angiopoietin-2 Quantikine ELISA Kit (R&D, MANG20). TNYL-RAW-Fc was measured as described (11). Death-related proteins were measured in cell lysates by a solid-phase antibody array with chemiluminescent detection (R&D Systems, 893900).

### Immunoprecipitation (IP) and Western blotting

IP and Western blotting were performed essentially as described (10,11); details and list of antibodies used are in Supplementary Materials and Methods.

### Study approvals and tissue specimens

All animal experiments were approved by the Institutional Animal Care and Use Committee of the CCR, NCI, NIH and conducted in adherence to the NIH Guide for the Care and Use of Laboratory Animals. All human cancer tissue specimens were obtained through approved protocols of the NCI/CCR or through a CCR-approved MTA (Cureline Inc.). Tissue specimens of mouse melanoma from the M1 and M2 models (12), were gifts of Drs. A. Sassano, C-P. Day, E. Perez-Guijarro and G. Merlino (CCR/NCI/NIH).

### Animal experiments

Female and male 8-week old C57BL/6J mice (Jackson Laboratories, 000664); female 10-week old BALB/cJ mice (Jackson Laboratories, 000651); female 8-week-old NOD-*scid*IL2Rgamma^null^ mice (Jackson Laboratories, NSG™; 005557), and female 6-10 week old Nu/Nu mice (6-10 week old Charles River Laboratories) were inoculated subcutaneously (s.c.) with B16F10, 4T1, LLC1, 9013BL/M2, Mel114433/M1 and MOPC-315 (1.0-2.0 ×10^6^cells). Tumor volume was estimated by V=D(d^2^)/2 (D longest and d shortest perpendicular dimensions). Details of animal experiments are found in Supplementary Materials and Methods. Briefly, groups of mice were randomized to receive: daily administration via gavage of 0.1 ml formulation buffer or SHP099 (80-200 mg/kg) in 0.1 ml formulation buffer; twice/week s.c. injections of PBS alone (0.1ml) or AMG386 (5.6 mg/kg in 0.1 ml PBS); oral daily administration of 0.1 ml formulation buffer + 0.1 ml s.c. PBS twice/week or SHP099 (100-200 mg/kg) in 0.1 ml formulation buffer plus AMG386 (5.6 mg/kg in 0.1 ml PBS) twice/week. All mice were euthanized when tumor(s) in any mouse from any experimental group reached a size of 20 mm in any direction. TNYL-RAW expression in mice was achieved as described (11). Blood perfusion and tissue hypoxia were measured as detailed in Supplementary Materials and Methods.

### Immunofluorescence, imaging and image quantification

Tissue samples were fixed with cold 4% PFA over 72 hours at 4°C and processed as described (10,11). List of antibodies, other reagents and methods for immunostaining are detailed in Supplementary Materials Methods. For imaging, extended field of view tile image of tissue sections were acquired using a Zeiss LSM780 laser scanning confocal microscope (Carl Zeiss, Oberkochen, Germany) equipped with a 20× plan-apochromat (N.A. 0.8) objective lens and 32-channel GaAsP spectral detector. Details of image acquisition, processing and analysis are found in Supplementary Materials and Methods.

### Data analysis and statistics

Publicly available datasets are from The Cancer Gene Atlas (TCGA), NCI Genomic Data Commons (GDC) Data Portal (RNA-Seq for colon cancer and HTseq counts for melanoma) and NCBI Gene Expression Omnibus (GSE7553 https://pubmed.ncbi.nlm.nih.gov/18442402/). Mann–Whitney U‐test for comparison of gene expression between tumor and control and Kaplan–Meier survival analysis were performed using statistical computing and graphics software R (version 3.5.2). Results are presented as mean±S.D. No sample was excluded from analysis and no sample was measured repeatedly. The statistical significance of differences between two groups was calculated using two-tailed Student’s t-test. Statistical significance among multiple groups was calculated by ANOVA with Dunnett’s multiple comparison test. P values <0.05 were considered statistically significant.

## Results

### Identification of a mouse model for EphrinB2/SHP2 targeting in the tumor vasculature

Based on our previous experiments showing that tyrosine phosphorylated (p)-EphrinB2/SHP2 signaling delays the physiological involution of hyaloid vessels by blocking pro-apoptotic STAT signaling, we now examined if EphrinB is active in the tumor vasculature and contributes to preserve the viability of tumor vessels. To this end, we first searched for an ideal tumor model in the immunocompetent mouse in which the tumor vasculature is broadly p-EphrinB-positive, but the tumor cells and other tumor-infiltrating cells are p-EphrinB-negative. Such model would aid in dissecting the specific role of active EphrinB2 in the tumor vasculature.

Among the six mouse models analyzed, we found that p-EphrinB is present in most CD31^+^ vessels of B16F10 melanoma, whereas the tumor cells, pericytes, and other cells in the tumor microenvironment are p-EphrinB-negative (Fig.1A-C). B16F10 cells are also EphrinB2-negative (Fig. 1D). The other six mouse cancer models differed from the B16F10 model in having only a proportion of vessels displaying p-EphrinB (Lewis lung carcinoma/LLC1, 9013BL/M2 and Mel114433/M1 melanoma (12), HT29 colon, MOPC-315 plasmacytoma and displaying p-EphrinB2 in infiltrating Gr1^+^ myeloid cells (mammary 4T1) (Supplementary Fig. 1A,B and (13)). In addition, while B16F10 tumor cells are p-EphrinB-negative, the HT29 and to a lower degree 4T1, 9013BL/M2 and Mel114433/M1 cancer cells are p-EphrinB-positive (Supplementary Fig. 1A and (13)). Thus, among the six models analyzed, the B16F10 mouse tumor model fulfills the criteria of displaying broadly active EphrinB in the vascular endothelial cells but not in the tumor cells and other infiltrating cells.

**Figure 1.**
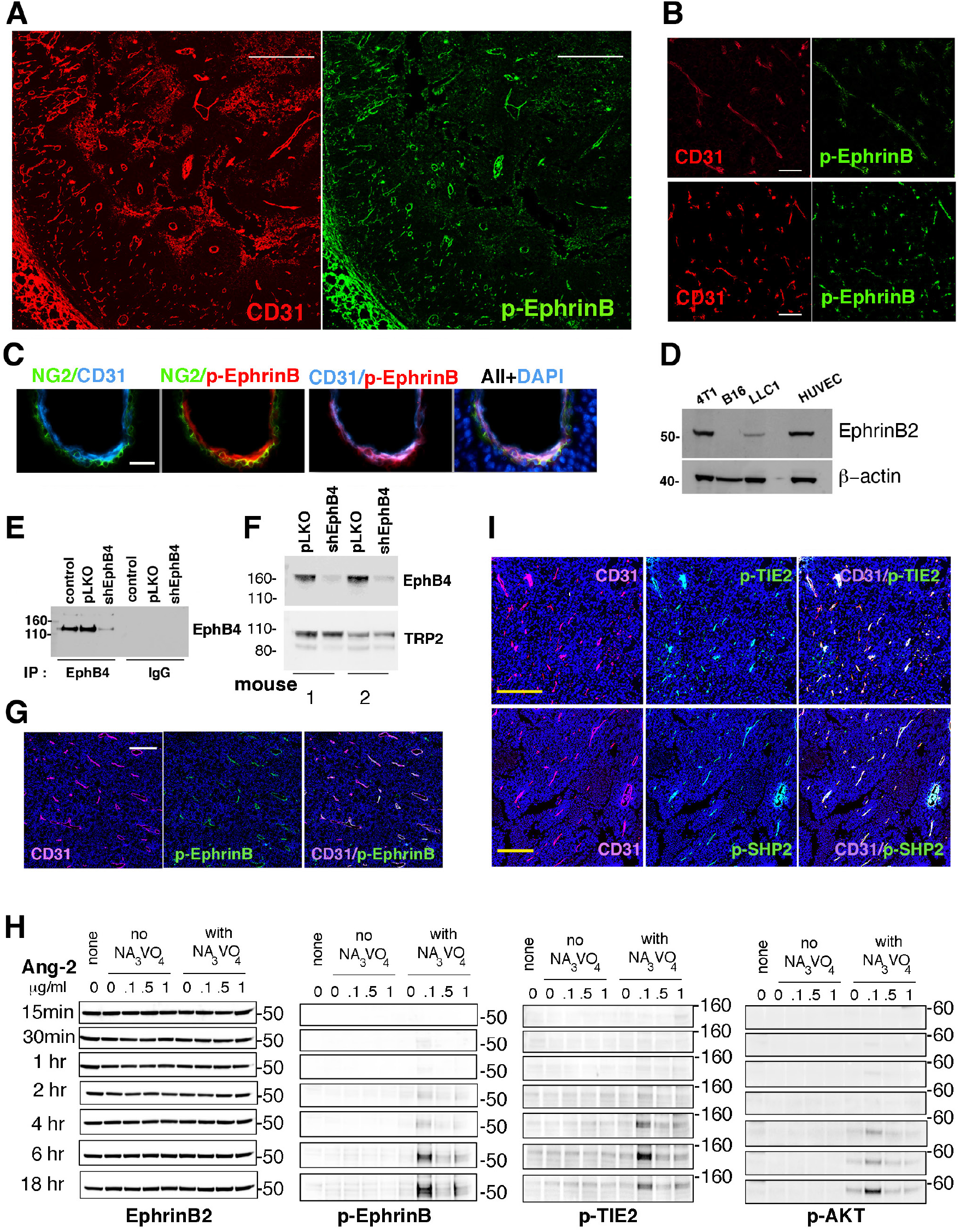
EphrinB2, TIE-2 and SHP2 phosphorylation in mouse tumor vessels. **(A,B)** Representative B16F10 tumor established in a syngeneic mouse showing that tumor vessels, identified by CD31 immunostaining, are generally p-EphrinB^+^. **(C)** the CD31^+^ endothelial cells lining a vessel are p-EphrinB^+^ whereas NG2^+^ perivascular cells are p-EphrinB^-^; cell nuclei are identified by DAPI; confocal images (scale bars: 500μm in A; 50μm in B; 10μm in C). **(D)** EphrinB2 is detected in cell lysates of 4T1, LLC1 and HUVEC, but not in B16F10 cells; immunoblotting. **(E)** EphB4 depletion in B16F10 cells; immunoprecipitation/immunoblotting; Control: no infection; pLKO: infection with control lentivirus; shEphB4: infection with silencing lentivirus. **(F)** B16F10 tumors induced by inoculation of EphB4-depleted B16F10 cells contain reduced levels of EphB4 compared to control pLKO-infected B16F10; immunoblotting; TRP2: tyrosine-related protein 2 (TRP2). **(G)** CD31^+^ vessels in EphB4-depleted tumors are generally p-EphrinB^+^; scale bar: 200mm. **(H)** Ang-2 dose and time-dependently activates p-EphrinB, p-TIE2 and p-AKT in HUVEC; immunoblotting results; NA3VO4: sodium orthovanadate. **(I)** Tumor vessels in B16F10 tumors are p-TIE2 (Tyr^992^)^+^ and pSHP2(Tyr^542^)^+^; confocal images (scale bars: 200μm).

EphrinB2 is normally activated by receptor engagement, particularly by EphB4 (8). We previously determined that EphB4 protein is present in B16F10 tumor cells (11). To test if B16F10-associated EphB4 activates tumor vascular EphrinB2, we silenced EphB4 in B16F10 cells (Fig. 1E) and generated syngeneic B16F10 tumors depleted of EphB4 (Fig. 1F). The vasculature within these EphB4-negative tumors was p-EphrinB^+^ (Fig. 1G). In addition, robust expression of the EphB4/EphrinB2 blocking TNYL-RAW peptide (11) did not reduce vascular p-EphrinB in B16F10 tumor-bearing mice (Supplementary Fig. 1C,D). Among the other EphB receptors that can activate EphrinB2, B16F10 expresses low levels of EphB6 and EphA4 (Supplementary Fig. 2A) whereas EphB1, EphB2 and EphB3 mRNAs are undetected (not shown). These experiments indicated that EphBs in B16F10 cells are unlikely responsible for EphrinB activation in the B16F10 tumor vasculature.

**Figure 2.**
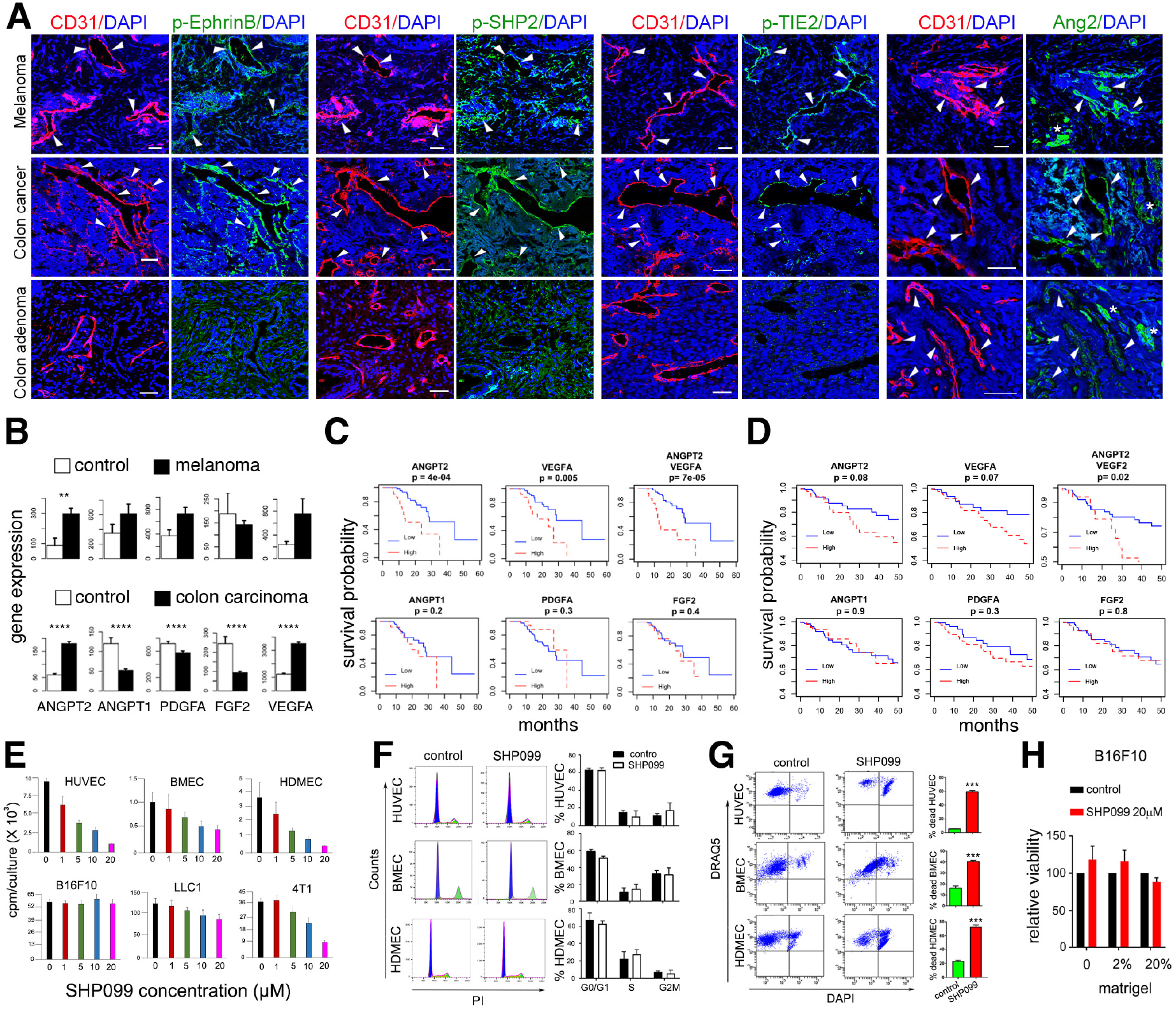
EphrinB2, TIE-2 and SHP2 phosphorylation in human melanoma and colon carcinoma. SHP2 inhibition in endothelial cells. **(A)** Representative tissue immunostaining of human melanoma, colon carcinoma and colon adenoma; confocal images (scale bars: 50μm tissues). **(B)** Gene expression in tumor (black bar) and control (white bar); top panel: melanoma (GSE7553, primary and metastatic; tumor n=54; control skin n=5); bottom panel: colon cancer (TCGA, primary and metastatic; tumor n=269; control n=41. **(C)** Impact of high and low (75% quartile) mRNA levels on the survival probability of patients with primary melanoma; Kaplan-Meier analysis using TCGA primary melanoma datasets (n=103); Mann Whitney U-test; \*\**P*<0.01 \*\*\*\**P*<0.0001. **(D)** Impact of high and low (median) mRNA levels on the survival probability of patients with colon cancer; Kaplan-Meier analysis using TCGA colon cancer dataset (n=262). **(E)** Proliferation of endothelial (HUVEC, BMEC and HDMED) and tumor (B16F10, LLC1 and 4T1) cells in three-day cultures supplemented with SHP099. Representative experiments of 3-8; error bars: ± S.D.; 5-10 replicate cultures/experiment. **(F)** Representative cell cycle flow cytometry profiles (left) and quantification of triplicate experiments (right); error bars: ± s.d. **(G)** Non-viable endothelial cells identified by flow cytometry (DRAQ5^+^/DAPI^+^) after 3-day culture with or without SHP099 (5μm HUVEC and HDMEC; 20μm BMEC); error bars: ± s.d.; triplicate experiments. **(H)** Effect of Matrigel on B16F10 cell viability. Representative experiments of 3; 5 replicate cultures; error bars: ± S.D.

In addition to becoming phosphorylated through binding interaction with EphB receptors and subsequent Src kinases recruitment, the EphrinB1 intracellular domain is trans-phosphorylated in cis by certain TK growth factors receptors, including TIE-2, FGFR and PDGFR (8,9,14). We found that B16F10 cells express higher mRNA levels of the TIE-2 ligand Angiopoietin-2 (Ang2) compared to LLC1 and 4T1 tumor cells, whereas Ang1, Platelet-Derived Growth Factor Alpha (PDGFA) and Fibroblast Growth Factor2 (FGF2) are similarly expressed (Supplementary Fig. 2B). B16F10 cells also secrete Ang2 in the culture supernatant (Supplementary Fig. 2C) and B16F10 tumor-bearing mice have significantly more circulating Ang2 than control mice (Supplementary Fig. 2D). In addition, Ang2 is detected in B16F10 tumor tissues, particularly in the cytoplasm of the tumor cells (Supplementary Fig. 2E).

We therefore tested whether Ang2 can activate EphrinB2 in endothelial cells through its TIE2 TK receptor (15). Recombinant Ang2 dose and time-dependently induced TIE2^Y992^, EphrinB2^Y324/329^ and AKT^Ser473^ phosphorylation in primary human umbilical vein endothelial cells (HUVEC, Fig. 1H). These results suggested that p-TIE2 may contribute to activation of EphrinB2 in the tumor vasculature of B16F10 tumors. Consistent with this possibility, we found that TIE-2 is broadly tyrosine phosphorylated in the vasculature of B16F10 tumors (Fig. 1I).Additionally, we found that the CD31^+^ endothelial cells of B16F10 tumors are generally p-SHP2^Tyr542^-positive, whereas the tumor cells are largely p-SHP2^-^ (Fig. 1I). These results indicated that the B16F10 mouse tumor model provides an opportunity for selectively targeting p-EphrinB/SHP2 in the tumor vasculature and to investigate relationships between p-EphrinB/SHP2 and Ang2/TIE2 signaling.

### EphrinB2/SHP2 and Ang2/TIE2 activity in vascular endothelial cells of human cancers

A screen of biopsy specimens from human melanoma and colon carcinoma showed that most CD31^+^ endothelial cells in these tissues are p-EphrinB^+^, pSHP2^+^ and p-TIE2^+^ (Fig. 2A) and that Ang2 is present both in CD31^-^ and CD31^+^ cells in the tumor tissue (Fig. 2A). In human colon adenomas, however, the endothelial cells are mostly p-EphrinB^-^, pSHP2^-^, pTIE2^-^ and Ang2^+^ whereas the CD31^-^ cell are mostly Ang2^-^ (Fig. 2A). Specimens from human breast adenocarcinoma, lung carcinoma, glioblastoma, angiosarcoma and Kaposi’s sarcoma showed the presence of p-EphrinB, p-SHP2 and p-TIE2 in a proportion of tumor vascular endothelial cells and Ang2 in a proportion of the CD31^-^ tumor cells (not shown). Our detection of Ang2 in these cancers is consistent with previous observations showing that Ang2 is expressed not only in endothelial cells but also in some cancer cells (16).

Since these results indicate that the EphrinB2/SHP2 and Ang2/TIE2 pathways are active in the tumor vasculature of human melanoma and colon carcinoma, we examined the potential impact of vascular EphrinB2/SHP2 and Ang2/TIE2 activity on disease outcome. We focused on Ang2 expression in human melanoma and colon carcinoma, since Ang2 drives both TIE2 and EphrinB activation in endothelial cells. In addition, since active PDGFR and FGFR can trans-phosphorylate EphrinB2, we also examined expression of their specific ligands PDGFA and FGF2, along with Ang1 and VEGFA in these cancer tissues.

*ANGPT2* gene expression is significantly higher (P<0.01) in skin melanoma compared to normal skin, whereas expression of *ANGPT1, PDGFA, FGF2* and *VEGFA* is not significantly different (Fig. 2B). In addition, high *ANGPT2* or high *VEGFA* expression in primary melanoma predicts a lower probability of survival compared to low *ANGPT2* (P=0.0004) or *VEGFA* (P=0.0052) expression. Together, high *ANGPT2* plus high *VEGFA* expression also predicts a significantly reduced probability (P=0.0002) of survival compared to low Ang2 plus low *VEGFA* expression (Fig. 2C). However, high or low expression of *ANGPT1, PDGFA* and *FGF2* has no impact on the survival probability of patients with primary melanoma.

In colon carcinoma, *ANGPT2* and *VEGFA* expression is significantly higher (P<0.0001) in the tumor compared to the adjacent normal colon, whereas expression of *ANGP1, PDGFA* and *FGF2* is significantly lower (P<0.0001) (Fig. 2B). In patients with colon carcinoma (n=262), high *ANGP2* and *VEGF2* expression, individually, predicts a somewhat worse probability of survival compared to low *ANGPT2* or *VEGFA* expression, whereas high or low *ANGPT1, PDGFA* and *FGF2* expression has no impact (Fig. 2D). Together, high A*NGPT2* plus high *VEGFA* expression predicts a significantly (P=0.02) lower probability of survival in patients with colon carcinoma (Fig. 2D). Similar conclusions on the impact of *ANGPT2* and *VEGFA* expression on probability of colon cancer survival emerges from analysis of an additional colon carcinoma dataset (not shown, Affymetrix microarray E-17538).

These results indicate that the EphrinB/SHP2 and Ang2/TIE2 pathways are active in the vasculature of human melanoma and colon carcinoma, and that high expression of ANGPT2 alone or with high VEGFA in these cancers predicts a worse patients survival probability.

### Inhibition of SHP2 tyrosine phosphatase compromises endothelial cell viability

To evaluate the impact of SHP2-regulated signaling on endothelial cell survival, we utilized the allosteric SHP2 inhibitor SHP099, which stabilizes SHP2 in an auto-inhibited conformation and renders it enzymatically inactive (17,18). SHP099 dose-dependently reduces the proliferation of HUVEC, mouse bone marrow endothelial cells (BMEC) and human dermal microvascular endothelial cells (HDMEC); it also inhibits proliferation of the murine 4T1 and LLC1 cancer lines (Fig. 2E). However, SHP099 (1-20 μm) has minimal effect on the proliferation of murine melanoma B16F10 cells (Fig. 2E). SHP099 inhibits endothelial cell proliferation after 24-48 hours incubation (Supplementary Fig. 3A) with no or minimal change in cell cycle distribution (Fig. 2F). After 3-day incubation, SHP099 induces cell death in HUVEC (5μm), HDMEC (5μm) and BMEC (20μm) (Fig. 2G); residual endothelial cells resume growth after wash-removal of SHP099 with kinetics comparable to control cells (Supplementary Fig. 3B). However, SHP099 does not reduce the viability of B16F10 cells even when the cells are cultured in 2-20% Matrigel, under conditions that have revealed increased sensitivity to SHP099 in certain KRAS-mutant cancer cells (19) (Fig. 2H).

**Figure 3.**
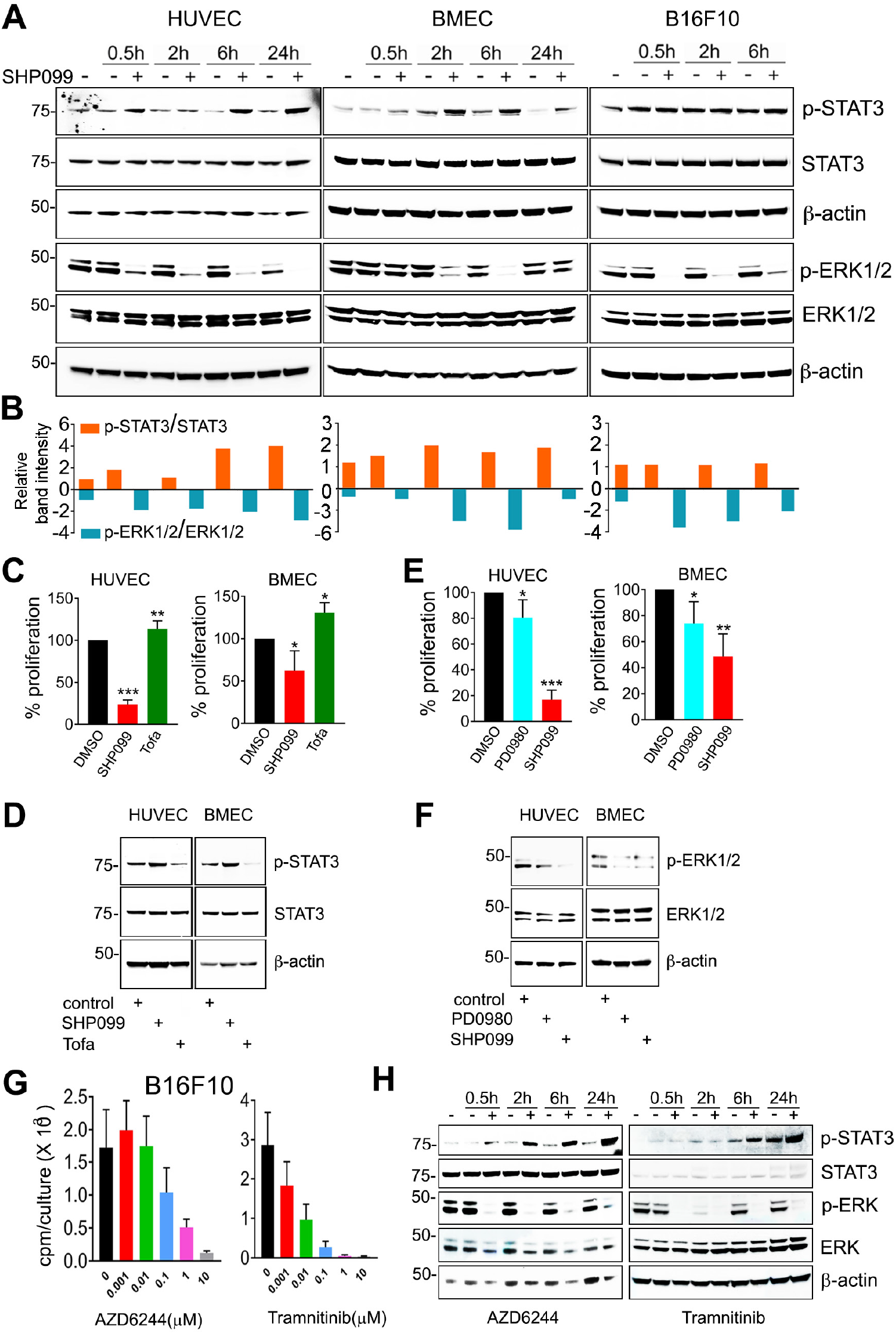
SHP099 reduces ERK1/2 and increases STAT3 phosphorylation in endothelial cells. **(A)** Kinetic SHP099 modulation of p-STAT3 (Tyr^705^) and p-ERK1/2 (Thr^202^/Tyr^204^) in endothelial and B16F10 cells; SHP099: 5μm HUVEC; 20μm BMEC and B16F10; representative immunoblotting of 3-5 experiments; h: hours. **(B)** Quantification of band intensities in panel (a); p-STAT3/STAT3 and p-ERK1/2/ERK1/2. **(C,D)** The JAK inhibitor Tofacitinib (Tofa, 50nM) enhances the proliferation of HUVEC and BMEC compared to control and reduces endogenous p-STAT3 levels in HUVEC and BMEC compared to control (D) after 72-hour culture. SHP099 (5μm HUVEC; 20μm BMEC) reduces cell proliferation and increases p-STAT3 in HUVEC and BMEC compared to control. **(E,F)** The MAP kinases inhibitor PD098059 (PD0980, 10μm) and SHP099 (5μm HUVEC; 20μm BMEC) inhibit HUVEC and BMEC proliferation compared to control (E) and reduce p-ERK1/2 levels by immunoblotting after 72-hour culture (F). Results of proliferation (% of control) from triplicate cultures are representative of 3-4 experiments; \**P*<0.05, \*\**P*<0.01 \*\*\**P*<0.001; two-tailed Student’s t-test; error bars: ± S.D. **(G,F)** The ERK1/2 inhibitors AZD6244 and Tamnitinib dose-dependently reduce B16F10 cell proliferation (G); and levels of endogenous p-STAT3 and p-ERK in B16F10 cells (H).

In endothelial cells, SHP099 induces the accumulation of cell-death-associated proteins, including cleaved-caspase-3, and the reduction of pro-survival proteins, including Survivin (Supplementary Fig. 3C). Also, SHP099 reduced endothelial cell content of vascular endothelial (VE)-cadherin (Supplementary Fig. 3D), an endothelial pro-survival protein (20) identified as a mediator of SHP2 protective function in endothelial cells (21). Furthermore, SHP2 increased the levels of the pro-apoptotic JNK3 in BMEC and HUVEC (Supplementary Fig. 3E). These results show that SHP099 reduces endothelial cell proliferation and induces cell death in endothelial cells. Consistent with these results, depletion of SHP2 from primary endothelial cells was reported to reduce cell proliferation, induce cell death (22), and destabilize endothelial cell junctions reducing endothelial cell barrier functions (21).

### SHP2 modulates STAT and MAPK signaling in endothelial cells

The SHP2 phosphatase has many substrates and has been linked to regulation of several signaling cascades, including the ERK/MAPK and STAT cascades (23,24). To identify the mechanisms underlying SHP099 inhibition of endothelial cell survival, we first focused on STAT signaling since we previously found that SHP2 silencing activates this pathway in endothelial cells (10). We now observed that SHP099 activates STAT3 in HUVEC (5μM) and BMEC (20μM), but not in the melanoma B16F10 cell line (20μM) (Fig. 3A,B). At these concentrations, SHP099 did not activate STAT1 or STAT5 and did not significantly alter AKT activity in endothelial and B16F10 cells (Supplementary Fig. 4A-E).

**Figure 4.**
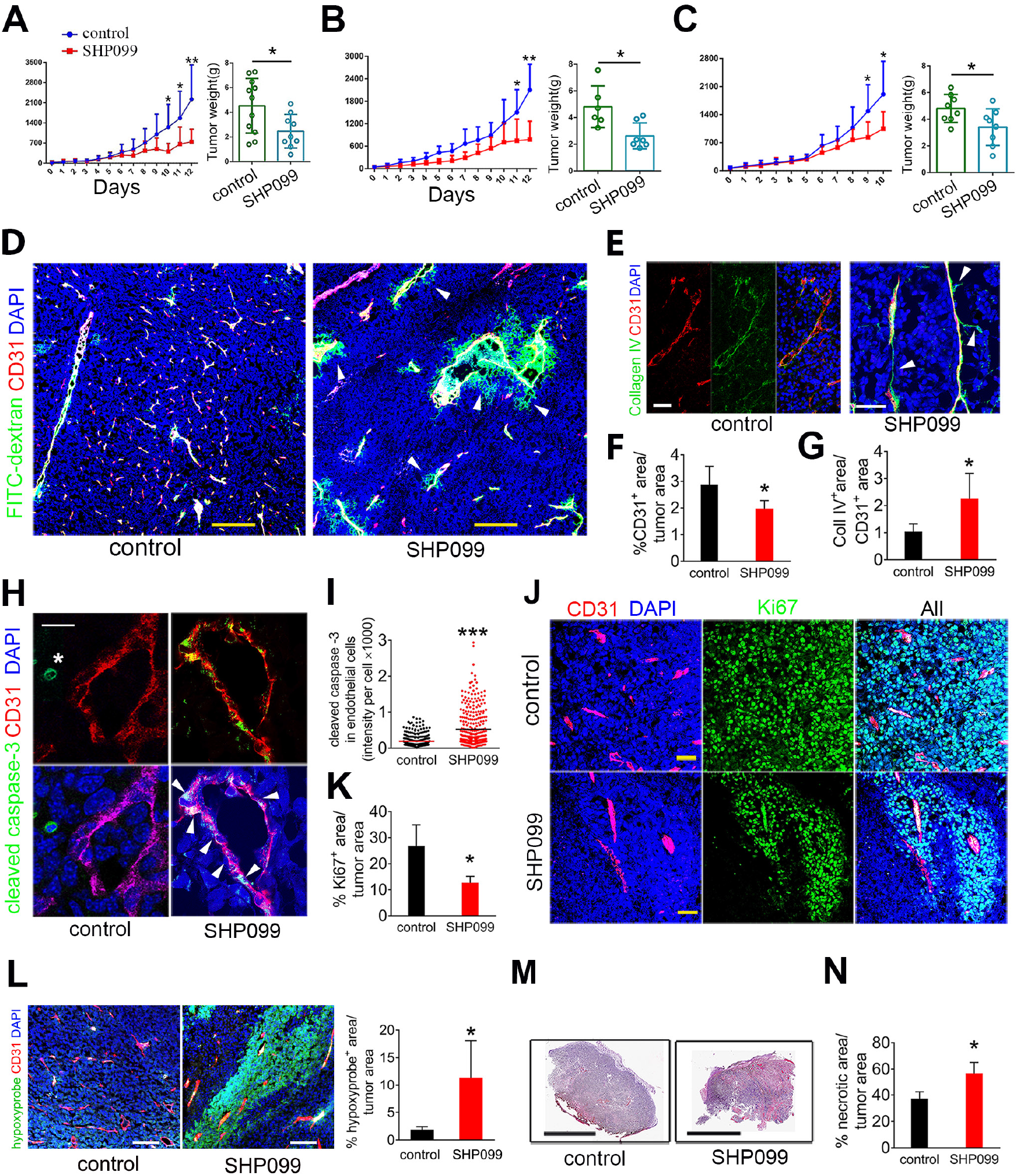
Vascular effects of SHP099 and inhibition of tumor growth. **(A-C)** SHP099 (A, 75mg/kg; B, C: 100mg/kg) reduces B16F10 tumor growth in syngeneic mice. Panels depict tumor growth from initiation of treatment to endpoint (left) and tumor weight (right) at endpoint; no. mice/group: 6-11 control; 7-9 SHP099. \**P*<0.05 and \*\**P*<0.01 by two-tailed Student’s t-test. Extravasation of FITC-dextran (green, pointed by arrowheads) in tumors treated with SHP099. Vessels identified by CD31 (red); DAPI (blue) identifies cell nuclei. Confocal images (scale bar: 200μM). **(E)** “Vascular sleeves” (Collagen IV^+^ CD31^-^) identified in a representative SHP099-treated tumor. Confocal images (scale bar: 50μM). **(F)** Reduced CD31^+^ vascular area in SHP099-treated tumors compared to controls (n=6/group); quantification by ImageJ. **(G)** The ratio of Collagen IV^+^/CD31^+^ area is increased in SHP099-treated tumors compared to controls (n=6/group); quantification by ImageJ; Coll: collagen. **(H)** Cleaved caspase-3 detection in a CD31^+^ vessel (representative SHP099-treated tumor); arrowheads point to cleaved caspase-3^+^CD31^+^ cells. Cleaved caspase-3^+^ tumor cell is marked by *. Confocal images (scale bar: 10μM). **(I)** Increased cleaved caspase-3 fluorescence intensity in CD31^+^ cells from SHP099-treated tumors compared to controls (n=3/group); at least 100 cells counted/sample. **(J)** Ki67^+^ proliferating cells in representative SHP099-treated and control tumors; CD31 identifies the vessels; DAPI identifies cell nuclei. Confocal images (scale bar: 50μM). **(K)** Quantification of proliferating cells within SHP099 and control (n=4/group) tumors by ImageJ. **(L)** Hypoxic tumor identified Hypoxyprobe (green); confocal images (scale bar: 100μM), left; quantification of hypoxic areas in control and SHP099-treated tumors by ImageJ. **(M)** Representative control and SHP099-treated B16F10 tumor sections through the maximum diameter; H&E staining (scale bar: 10μM). **(N)** Quantification of necrotic tumor areas by ImageJ in control and SHP099-treated tumors (n=3/group). Error bars: ± S.D.: P values from two-tailed Student’s t-test; \**P*<0.05, \*\**P*<0.01 and \*\*\**P*<0.001.

SHP2 is required for optimal activation of the ERK-MAP kinase pathway that sustains cell survival and proliferation in many cell types (24), including endothelial cells (18,25). We found that SHP099 inhibits ERK activity in HUVEC, BMEC and B16F10 cells (Fig. 3A,B). Noteworthy, SHP099 reduces ERK activity, without activating STAT3 in B16F10 cells (Fig. 3A,B). In addition, SHP099 activates STAT3 and inhibits ERK activity in endothelial cells beginning at similar early time-points, suggesting that SHP2 may regulate these two pathways directly rather than change in one pathway be compensatory to the other.

Next, we evaluated the effects of STAT3 activation and ERK1/2 inactivation on endothelial cell proliferation. The JAK kinases inhibitor Tofacitinib (Tofa, 50nM) reduced endogenous p-STAT3 activity and enhanced the proliferation of HUVEC and BMEC compared to control (Fig. 3 C,D). In addition, the ERK1/2 inhibitor PD098059 (PD0980, 10μM) reduced endogenous p-ERK1/2 activity and inhibited the proliferation of HUVEC and BMEC compared to control (Fig. 3 E,F). Furthermore, Tofa increased and PD0980 diminished cell survival in BMEC (Supplementary Fig. 4F).

In a panel of cancer cell lines (Supplementary Fig. 4G), SHP099 reduced the proliferation of all but two cell lines (colon carcinoma RKO and MC38). In the growth-inhibited cell lines, SHP099 enhanced STAT3 and reduced ERK1/2 activity. However, in the RKO and MC38 lines, SHP099 neither reduced cell proliferation nor altered STAT3 and ERK1/2 activity. Since in B16F10 cells, SHP099 had no effect on cell proliferation and STAT3 activity, but reduced ERK1/2 activity (Fig. 3A,B), we tested the effects of two well-established MAPK/ERK pathway inhibitors in B16F10 cells. AZD6244 and Tramnitinib dose-dependently inhibited B16F10 cell proliferation (Fig. 3G), reduced ERK1/2 activity and additionally activated STAT3 (Fig. 3H). MEK inhibitors were previously reported to induce “compensatory” STAT3 activation in other cancer cells (26). Since SHP099, differently from MEK inhibitors, failed to induce STAT3 activation in B16F10 cells (Fig. 3A) these results raise the possibility that STAT3 activation may be a critical contributor to growth inhibition in B16F10 tumor cells. Importantly, these results indicate that SHP099 reduces endothelial cell proliferation and viability associated with STAT3 activation and ERK1/2 inhibition, suggesting that SHP099 may represent a previously unidentified anti-vascular agent.

### SHP099 promotes tumor vessel regression and reduces tumor growth

We selected the syngeneic B16F10-melanoma mouse model to evaluate the anti-vascular effects of SHP099. In vitro, B16F10 cells are more resistant to SHP099 growth inhibition than primary HUVEC and HDMEC (Fig. 2E) and the likelihood of SHP099 having a direct anti-tumor effect appeared low since SHP099 transiently yielded maximal free plasma concentrations of >20μM at the effective oral dose of 100mg/kg (18). Also, the SHP099 target, SHP2, is phosphorylated in the CD31^+^ vascular endothelial cells of B16F10 tumors whereas the CD31^-^ tumor cells are generally p-SHP2^-^ (Fig. 1I).

We established subcutaneous B16F10 tumors in syngeneic mice. Groups of mice (7-11 mice/group) bearing similar size (range 30-50 μM^3^) tumors were treated daily with oral SHP099 (75-100 mg/kg) or vehicle only; the experiment was terminated when any tumor reached the maximum diameter of 20 μM in any direction. In three independent experiments, SHP099 reproducibly reduced B16F10 tumor growth and tumor weight compared to control (Fig. 4A-C), without causing loss of body weight (Supplementary Fig. 5A,B) or evidence of other toxicity. This anti-tumor effect of SHP099 was associated with vascular leakage, as the FITC-dextran tracer (2,000,000 kDa) was frequently found outside the vessel wall in the tumor parenchyma where auto-fluorescent erythrocytes were also found (Fig. 4D and Supplementary Fig. 5C). Type IV collagen immunostaining to visualize vascular basement membranes showed scattered remnants of vessels (“sleeves” (27)) that lacked endothelial CD31 coverage in SHP099-treated but generally not in control tumors (Fig. 4E-G and Supplementary Fig. 5D). Tumor vascularity was also reduced (Fig. 4F). Further supporting vessel involution, cleaved-caspase-3-positive endothelial cells decorated the wall of tumor vessels from the SHP099-treated more frequently than in control mice (Fig. 4H,I and Supplementary Fig. 5E). The Ki67^+^ cells were reduced in SHP099-treated tumors compared to controls, indicative of reduced tumor cell proliferation (Fig. 4 J,K and Supplementary Fig. 5F). In addition, tumor tissue hypoxia (Fig. 4L) and necrosis (Fig. 4 M,N) were increased in SHP099-treated tumors compared to controls.

**Figure 5.**
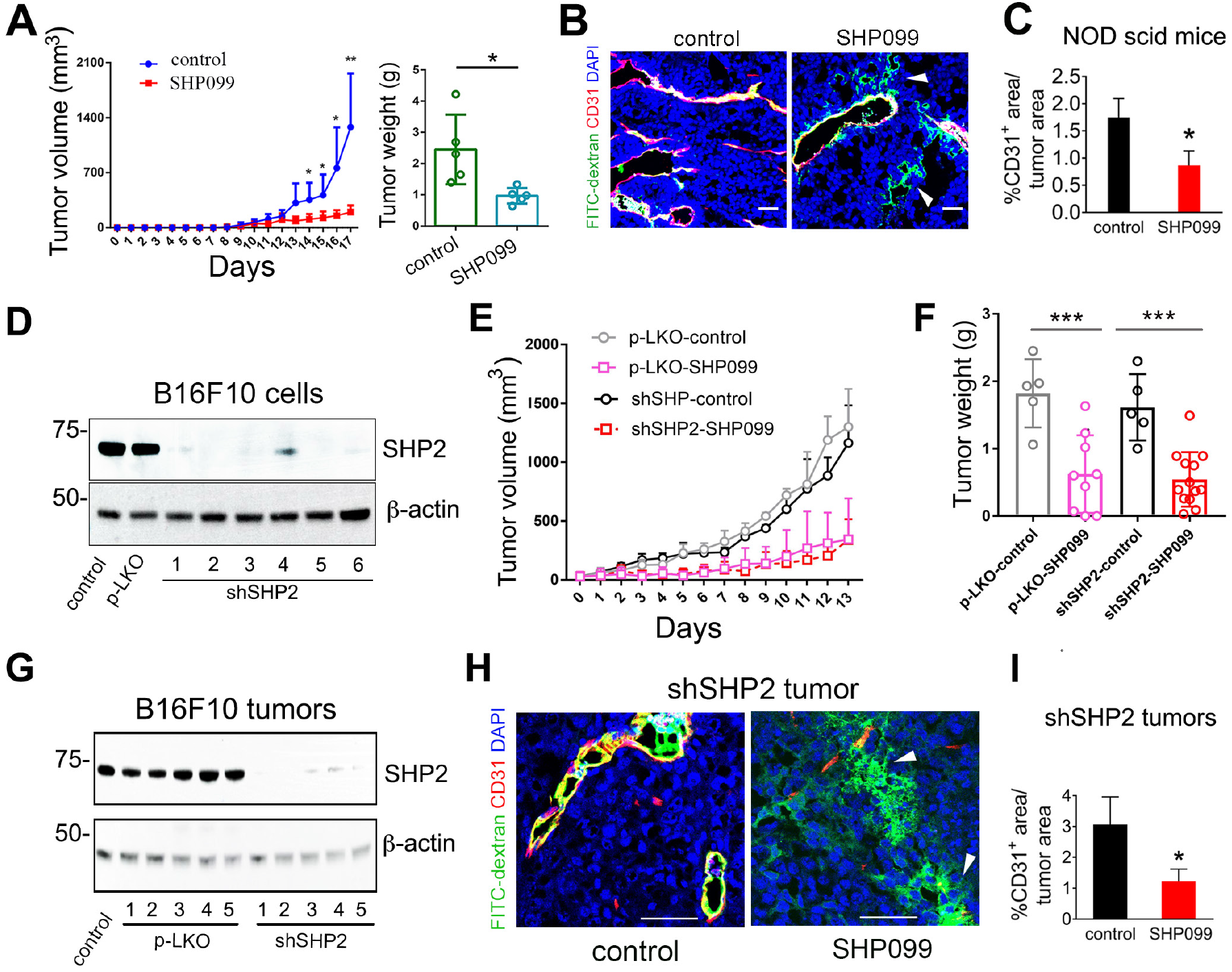
SHP099 inhibition of tumor growth in SHP2-depleted tumor cells and in immunodeficient mice. **(A)** SHP099 reduces the growth and weight of s.c. B16F10 tumors established in NOD scid (NSG) mice (n=5/group); P values from two-tailed Student’s t-test; \**P*<0.05, \*\**P*<0.01. **(B)** Extravasation of FITC-dextran (green) in tumors established in NOD scid mice treated with SHP099; scale bar (50μM). **(C)** Reduced CD31^+^ vascular area in SHP099-treated tumors compared to controls (n=5/group); quantification by ImageJ. P values from two-tailed Student’s t-test; \**P*<0.05. **(D)** Effective SHP2 depletion in B16F10 cells. Control: no infection; pLKO: infection with control lentivirus; shEphB4: infection with six silencing lentiviral preparations. **(E,F)** SHP099 reduces the growth (E) and weight (F) of established s.c. B16F10 tumors infected with control lentivirus (n=9) or with shSHP2-2 (n=13) compared to untreated controls; control lentivirus (n=5) and shSHP2-2 (n=12). P values by analysis of variance with Dunnett’s multiple comparison test; \*\*\**P*<0.001. **(G)** Persistent depletion of SHP2 in tumors induced by shSHP2-B16F10 cells (n=5) compared to control tumors induced by p-LKO-B16F10 (n=5); tumors were removed from mice 18 days after cell inoculation. **(H)** SHP099 induces extravasation of FITC-dextran (green) in tumors generated by inoculation of shSHP2-B16F10.; scale bar: 50μM. **(I)** Reduced CD31^+^ vascular area in SHP099-treated tumors induced by s.c. inoculation of shSHP2-B16F10 cells compared to controls induced by p-LKO-B16F10 cells (n=5/group); quantification by ImageJ. Error bars: ± S.D.: P values from two-tailed Student’s t-test; \**P*<0.05.

The prominent vascular pathology displayed by SHP099-treated tumors suggested a vascular basis of the anti-tumor effect of SHP099. The CD31^+^ vasculature in liver, lung, heart, intestine, brain and muscle tissues appeared normal and was normally perfused in tumor-bearing mice after SHP099 treatment, and the tissues appeared normal (Supplementary Fig. 6). Thus, the selectivity of SHP099 for the tumor vasculature may reflect differences between quiescent vessels in adult tissues and remodeling vessels of tumors.

**Figure 6.**
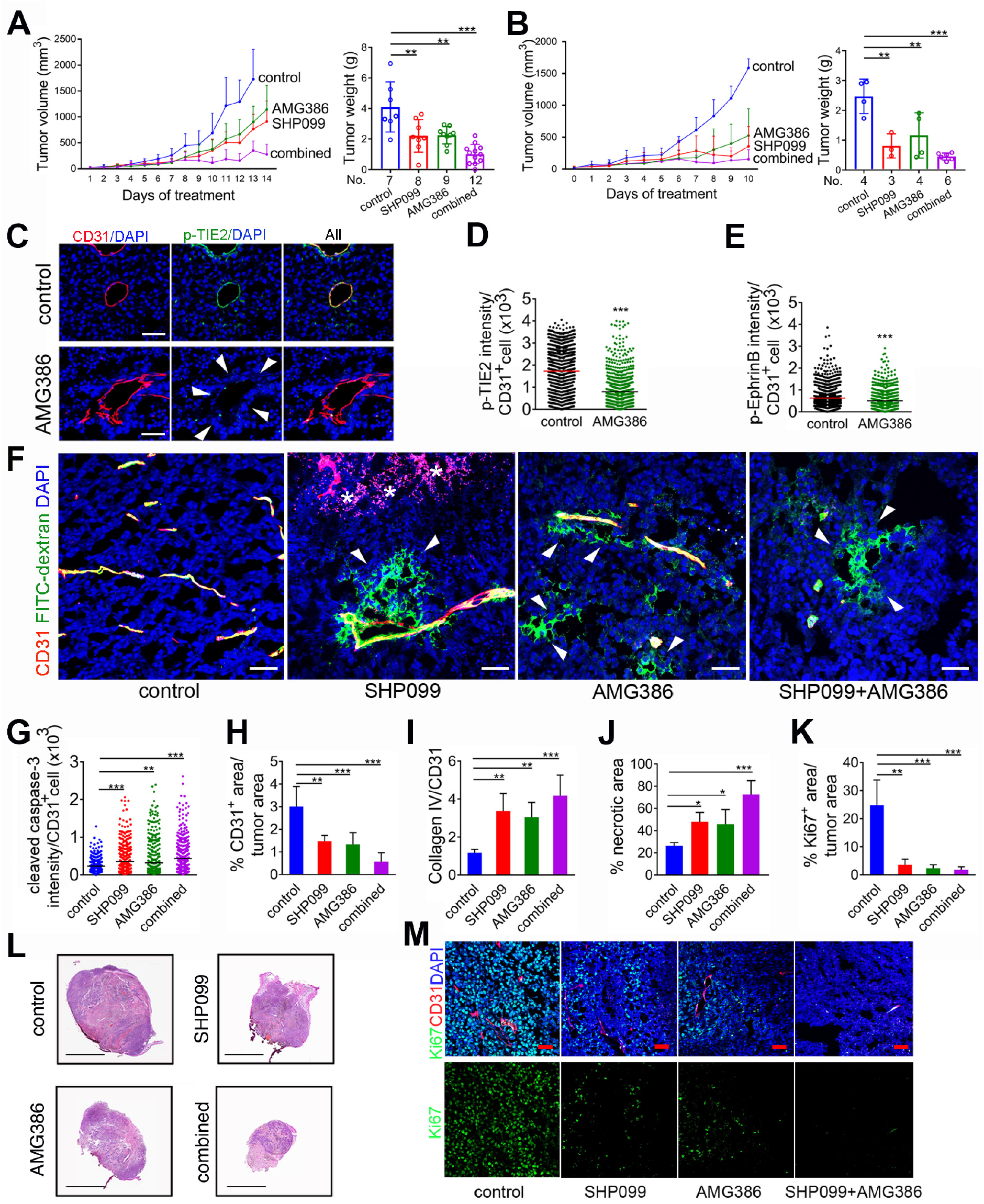
SHP099 and AMG386 cooperatively reduce tumor vascularization and growth. **(A,B)** AMG386 and SHP099 cooperatively reduce B16F10 tumor growth. AMG386 (5.6mg/kg s.c.; twice/week) and SHP099 (a: 200 mg/kg; b: 100mg/kg orally/daily) were administered individually or together to B16F10 tumor-bearing mice. Panels depict tumor growth from initiation of treatment to endpoint (left) and tumor weight (right) at endpoint; error bars ± s.d.; P values by analysis of variance with Dunnett’s multiple comparison test; \*\**P*<0.01, \*\*\**P*<0.001. **(C)** p-TIE2 in representative tumor vessels from control and AMG386-treated mice; confocal images (C), scale bar: 50μM). **(D,E)** Quantification of p-TIE2 (D) and p-EphrinB (E) in CD31^+^ tumor endothelial cells from control and AMG386-treated mice; P values from two-tailed Student’s t-test; \*\*\**P*<0.001. **(F)** Vessel perfusion visualized by FITC-dextran (green); CD31 (red) identifies the endothelium; DAPI (blue) identifies the nuclei; representative tumor confocal images (scale bar:50μM); arrow heads point to FITC-dextran extravasation; asterisks to extravasated erythrocytes. **(G)** Quantification of cleaved caspase-3 in the tumor vasculature; n=3/group; at least 100 CD31^+^ cells counted/sample. **(H)** Quantification of tumor vascular area; n=5/group. **(I)** Quantification of “vascular sleeves” (CD31^-^Collagen IV^+^) n=5/group. **(J)** % necrotic tumor area (n=4/group) and **(K)** % proliferating tumor cells (n=3/group) in control and treated groups. Error bars: ± s.d.; P values by analysis of variance with Dunnett’s multiple comparison test; \**P*<0.05, \*\**P*<0.01, \*\*\**P*<0.001. **(L)** Representative control and treated B16F10 tumor sections through the maximum diameter; H&E staining; (scale bar: 10μM). **(M)** Proliferating Ki67^+^ cells (green) in representative control and treated tumors; CD31 identifies the vessels; DAPI identifies cell nuclei. Confocal images (scale bar: 50μM).

SHP099 may elicit anti-tumor immune cell responses by attenuating signaling from immune-inhibitory receptors, such as PD-1 (18,28). We tested for a potential contribution of anti-tumor immunity to the anti-tumor activity of SHP099 in B16F10 tumors. Given that T and NK cells promote the effectiveness of checkpoint blockade (29,30), we utilized the T and NK-cell deficient NSG (NOD scid gamma) mice. Due to the rapid growth of B16F10 tumors in NSG mice, treatment was initiated 72 hours after tumor cell injection. SHP099 showed similar anti-tumor (Fig. 5A) and anti-vascular (Fig. 5B,C) activity in immunodeficient NSG and immunocompetent B6 mice. Also, SHP099 did not substantially change the number of F4/80^+^ tumor-infiltrating macrophages in these NSG mice (Supplementary Fig 7 A,B). These results suggest that SHP099 activation of anti-tumor immunity is not critical in the current system.

**Figure 7.**
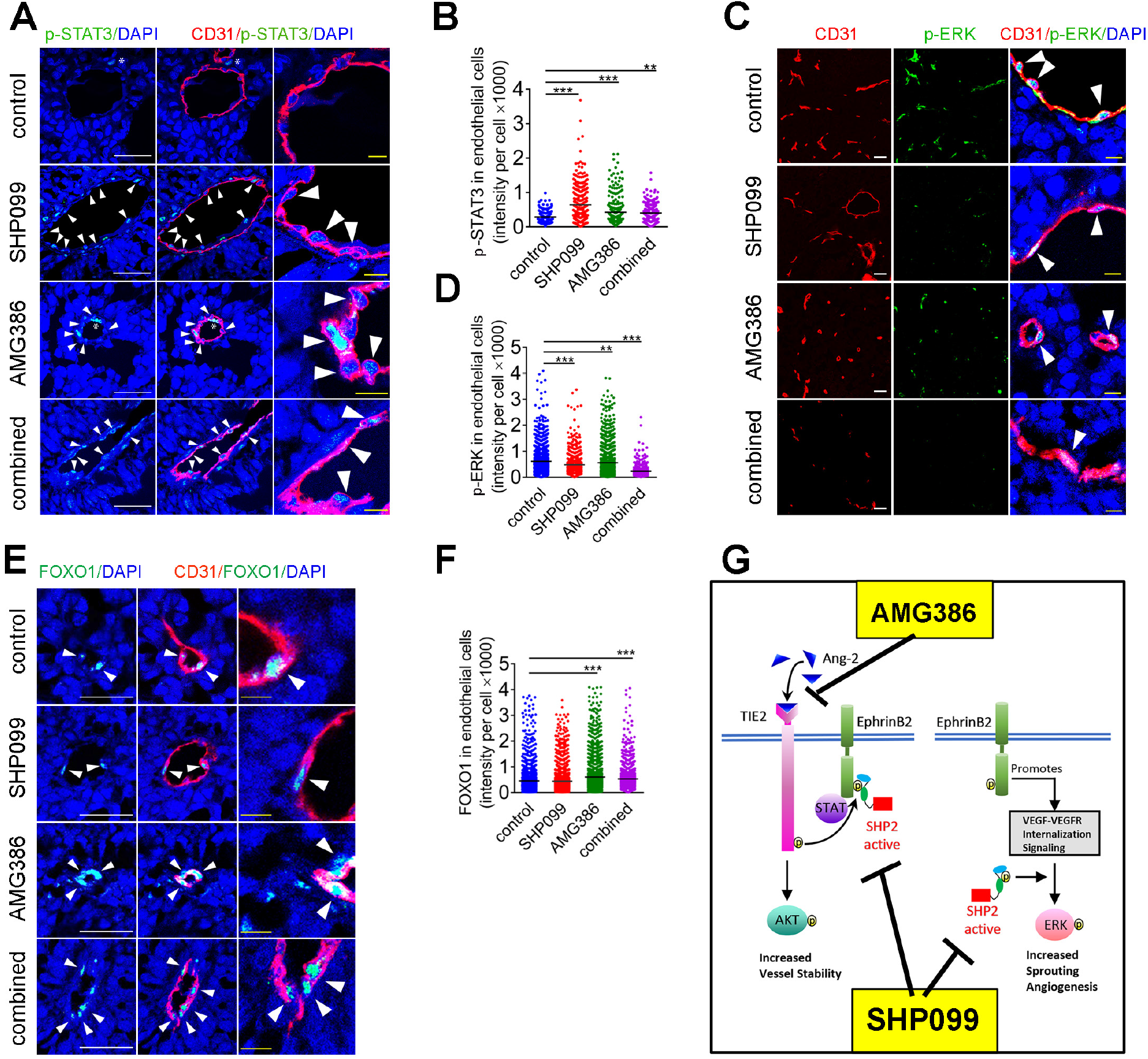
SHP099 and AMG386 modulate STAT3, ERK and AKT signaling in the tumor vasculature. Nuclear p-STAT3 **(A,B)** and p-ERK1/2 **(C,D)** in tumor CD31^+^ endothelial cells of mice treated for 14 days with SHP099 (200 mg/kg), AMG386 (5.6mg/kg) or the combination of SHP099 (200 mg/kg) + AMG386 (5.6mg/kg). Representative immunostaining (A,C) and quantification of staining intensity/CD31^+^ cell (B, D); horizontal line: mean intensity/group; n=3 tumors/group; at least 100 cells counted/sample. Scale bars: A, 50μM; C, 10μM. **(E,F)** FOXO1 detection in tumor CD31^+^ endothelial cells after treatment with SHP099 (200 mg/kg), AMG386 (5.6mg/kg) or the combination of SHP099 (200 mg/kg) + AMG386 (5.6mg/kg). Representative confocal images (E) and quantification (F) of FOXO1 fluorescence intensity/CD31^+^ cell. P values by analysis of variance with Dunnett’s multiple comparison test; \*\**P*<0.01, \*\*\**P*<0.001. **(G)** Schematic representation of the effects of SHP099 and AMG386 on signaling pathways active in the tumor vasculature.

SHP099 did not reduce the proliferation or viability of B16F10 cells *in vitro*. However, it remained possible that tumor cells acquired responsiveness to SHP099 *in vivo*, which could contribute to the anti-tumor activity of SHP099. To address this possibility, we stably depleted SHP2 from B16F10 cells using six different lentiviral silencing vectors (Fig. 5D) and injected SHP2-depleted (vector 2) B16F10 cells into syngeneic mice to generate tumors along with control B16F10 cells. SHP099 (100 mg/kg orally/daily) reduced the growth of SHP2-depleted and non-depleted B16 tumors to a similar degree (Fig. 5 E,F), We confirmed that tumor SHP2 was markedly reduced at completion of the experiment (Fig. 5G). The vascular phenotype of increased vascular leakage (Fig. 5H) and reduced vascularization (Fig. 5I) was similar in SHP2-depleted and non-depleted tumors treated with SHP099. These results demonstrate that SHP099 is a previously unrecognized anti-vascular drug that damages the tumor endothelium and likely reduces tumor growth through a vascular mechanism.

### SHP2 and Ang/TIE2 inhibitors cooperatively inhibit tumor vascularization and tumor growth

Despite its effectiveness, SHP099 did not induce a complete involution of the tumor vasculature and did not eradicate B16F10 tumors (Fig. 4A-C). To target the vessels that persisted after SHP099 treatment, we sought to inhibit Ang2/TIE2-induced tumor angiogenesis that is mediated by the PI3K/AKT pathway (31,32). SHP099 does not block PI3K/AKT signaling in endothelial cells (Supplementary Fig. 4C,D) and other cell types (Supplementary Fig. 4E) (33). This pathway is active in the vasculature of B16F10 tumors, since TIE2 is broadly phosphorylated in the endothelial cells (Fig. 1I) and the tumor cells produce abundant Ang2 (Supplementary Fig. 2C-E). In addition, TIE2 phosphorylation persists in the tumor vasculature of mice treated with SHP099 (Supplementary Fig. 7C).

AMG386, a peptibody that inhibits Ang1/Ang2-TIE2 binding and impairs tumor angiogenesis (34,35) significantly reduced the growth of established B16F10 tumors compared to control (Fig. 6A,B). The anti-tumor effect of AMG386 (5.6 mg/kg s.c.; twice/week) was similar to that of SHP099 (P=0.93, exp.1, panel A; P=0.50 exp. 2, panel B). Together, SHP099 and AMG386 cooperatively reduced tumor growth and weight (Fig. 6A; SHP099 vs SHP099+ANG386: *P*= 0.006 exp. 1, 0.07 exp. 2; AMG386 vs SHP099+AMG386: *P*=0.0002 exp. 1, *P*=0.049 exp. 2); one of the tumors underwent full regression. Like SHP099, AMG386 minimally affected the proliferation of B16F10 cells (Supplementary Fig. 7D), which do not express TIE-2 (36). Mouse body weight was similar after treatment with AMG386 alone or with SHP099 (Supplementary Fig. 5B).

As a single agent, AMG386 reduced tumor vascular p-TIE2 (Fig. 6 C,D) and p-EphrinB (Fig. 6E), which is consistent with the results in vitro showing that Ang2 activates p-TIE2 and p-EphrinB in primary endothelial cells (Fig. 1H). In addition, AMG386 compromised the integrity of tumor vessels as reflected by localized FITC-dextran extravasation into the tumor parenchyma (Fig. 6F) and the presence of cleaved caspase-3^+^/CD31^+^ cells lining blood vessels (Fig. 6G and Supplementary Fig. 7E). Compared to control, AMG386 reduced the CD31^+^ vessel area (Fig. 6H) and promoted formation of vascular “sleeves” (Fig. 6I and Supplementary Fig. 7F). AMG386 reduced viable and proliferative tumor burden (Fig. 6 J-M).

Dual treatment with SHP099 and AMG386 resulted in a remarkable reduction of tumor vascularity (Fig. 6H) and viable/proliferative tumor burden (Fig. 6 J-M). Some of the residual tumor vessels appeared damaged and leaky (Fig. 6F, Supplementary Fig. 7E) and not viable based on the presence of cleaved caspase 3^+^/CD31^+^ endothelial cells lining vessels (Fig. 6G and Supplementary Fig. 7E) and detection of vascular sleeves (Fig. 6I and Supplemental Fig. 7F). Unlike the anti-vascular effects of SHP099+AMG386 in the tumors, the vasculature in the liver, lung, heart, intestine, brain and muscle of mice treated with AMG386+SHP099 appeared normal and normally perfused (Supplementary Fig. 8).

We examined the activity of intracellular signaling targets of SHP099 and AMG386 in the tumor vasculature. Nuclear p-STAT3 was detected at significantly higher levels in the tumor vasculature of mice treated with SHP099, AMG386 or SHP099+AMG386 compared to control (Fig. 7 A,B), but the increase in p-STAT3 induced by AMG386 alone or with SHP099, was quantitatively lower than induced by SHP099 alone (*P*<0.001 both comparisons). Also, nuclear/active p-ERK1/2 was significantly reduced in the tumor vasculature of mice treated with SHP099, AMG386 or SHP099+AMG386 compared to control (Fig. 7 C,D). The reduction of p-ERK1/2 from AMG386 treatment was significantly lower than from SHP099 treatment (*P*<0.001), whereas the combination of SHP099 and AMG386 was more effective than SHP099 alone (*P*<0.001) (Fig. 7D). These results, showing that SHP099 activates p-STAT3 and reduces p-ERK1/2 in tumor endothelial cells of mice, mirror the effects of SHP099 in endothelial cells *in vitro* (Fig. 3 A,B). Additionally, the results showing that AMG386 reduces p-ERK1/2 and activates STAT3 in tumor endothelial cells are consistent with the reduced activity of EphrinB in tumor endothelial cells of mice treated with AMG386 (Fig. 6C). It was previously known that active EphrinB2 not only plays an important role in restraining STAT signaling in endothelial cells (10) but also promotes VEGF/VEGFR internalization/signaling resulting in ERK1/2 activation (37,38).

As an indicator of Ang1/2-TIE2/p-AKT signaling in the tumor vasculature, we examined the subcellular localization of the transcription factor Forkhead box protein O1 (FOXO1) (39). Active AKT leads to FOXO1 translocation from the nucleus to the cytoplasm where it is rapidly degraded(32,40), predicting that reduced AKT activity would lead to accumulation of FOXO1. By immunofluorescence, FOXO1 was mostly restricted to the DAPI^+^ nuclei of CD31^+^ tumor endothelial cells in control and treated mice (Fig. 7E), attributable to the rapid degradation of cytoplasmic FOXO1. Quantitatively, the tumor vasculature of AMG386-only or AMG386+SHP099-treated mice was significantly enriched with FOXO1 compared to control mice, whereas the tumor vasculature of SHP099-treated mice was not (*P*=0.38) (Fig. 7F). Together, these results support a model (Fig. 7G) by which SHP099 and AMG386 modulate signaling cascades in endothelial cells reducing tumor vascularization.

## Discussion

In many cancer types, SHP2 is hyperactive due to mutations of the Ptpn11 gene, which encodes SHP2, or to aberrant signaling from protein-tyrosine kinases, and functions as a proto-oncogene (41). Hence, SHP2 is a therapeutic target in cancer. The specific SHP2 inhibitor, SHP099, effectively suppressed the growth of receptor-tyrosine-kinase-driven cancers in mouse xenograft models (18) and prevented cancer resistance to inhibitors of MEK (33,42,43) and anaplastic lymphoma kinase (ALK) (44).

Here, using chemical inhibition of SHP2 and genetic approaches to deplete cellular SHP2, we show that the SHP2 phosphatase plays a key role as a mediator of tumor vessel persistence and identify SHP2 as a new pharmacological target for the successful anti-vascular therapy of cancer. SHP2 is a ubiquitously expressed intracellular tyrosine phosphatase that promotes ERK signaling induced by growth factors, cytokines and hormones and supports cell survival and proliferation (24,41). SHP099 promoted the involution of the remodeling vasculature of tumors and inhibited tumor neovascularization while having no detectable direct effect on the tumor cells or the resting vasculature of normal tissues, which substantially differs from the tumor vasculature (5). A recent study noted that vascularization was generally reduced in tumors treated with SHP099, but it remained unclear whether this was attributable to a primary effect on the tumor vessels or an indirect consequence of tumor cell targeting (33). In our study, the anti-vascular activity of SHP099 is unlikely a consequence of an effect on the tumor cells, since the genetic depletion of SHP2 from the B16F10 tumor cells had no impact on the anti-vascular and anti-tumor activities of SHP099. Thus, B16F10 cancer model differs from the mutant KRAS^G12C^ MIAPaCA-2 cancer model, in which the effectiveness of SHP099 was reported to be cancer cell intrinsic and the effects of SHP099 were recapitulated by the genetic depletion of SHP2 in the cancer cells (19). A contribution of T/NK-induced anti-cancer immunity appears unlikely in the B16F10 model, since SHP099 exerted similar anti-vascular and anti-tumor activities in immunocompetent and immunodeficient NSG mice. Mechanistically, SHP099 inhibited ERK1/2 phosphorylation and activated STAT3 phosphorylation in endothelial cells from culture and the tumor vasculature. This signaling regulation likely mediates the anti-vascular effects of SHP099, since ERK1/2 signaling is critical to endothelial cell sprouting and angiogenesis (25) and STAT signaling promotes endothelial cell death and vessel involution (10,45). Thus, the current results spotlight a spectrum of SHP2 functions in the vascular endothelium and spur clinical development of SHP099 and similar inhibitors as novel anti-vascular agents.

Genetic experiments in mice showed that SHP2 is required for the formation of a hierarchically organized vessel network in the mouse yolk sac, linking SHP2 to regulation of post-angiogenic/vasculogenic remodeling events (46). Similarly, the knockout of SHP2 in endothelial cells caused embryonic hemorrhage and death attributed to disruption of endothelial cell junctions (21). Experiments in vitro suggested that SHP2 regulates endothelial adhesive functions and migration stemming from SHP2 binding to the intracellular domain of endothelial platelet-derived growth factor receptor-b (PDGFR-b), VE-cadherin and PECAM-1 (47,48). The knockdown of SHP2 transiently reduced FGF2+VEGF-dependent endothelial cell viability *in vitro* and *ex vivo* (22), delayed vessel regeneration in a mouse skin wounding model (49) and increased endothelial monolayer permeability (21). Also, SHP2 promoted VEGF-dependent ERK1/2 signaling (50), consistent with the role of EphrinB2-SHP2 signaling as a key regulator of VEGFR2 internalization and ERK signaling (37,38).

VEGF is a principal inducer of neovascularization and vascular permeability (1). Anti-VEGF/VEGFR2 therapies are effective at reducing angiogenesis and tumor growth in mouse tumor models and in patients with certain cancers (1). The PLCγ-ERK1/2 cascade, which mediates VEGF/VEGFR2 signaling from phosphorylated tyrosine 1173, plays a central role in VEGF-induced proliferation/angiogenic sprouting (50). By inhibiting ERK1/2 signaling, SHP099 is expected to reduce VEGF-induced sprouting angiogenesis.

Consistent with responses to anti-VEGF therapy, SHP099 incompletely blocked B16F10 tumor neovascularization. Ang2, FGF2, IL-6, IL-10 and other factors have been linked to anti-VEGF resistance (1-3). Here, we identified Ang2/TIE2 as a pro-angiogenic pathway active in B16F10 tumors, which secrete Ang2 (16), and found that Ang1/2-TIE2 neutralization enhances the efficacy of SHP099 as an anti-angiogenic and anti-cancer agent. Consistent with this, inhibition of Ang2 contributed to overcome resistance to VEGF blockade (35). Ang1/TIE2 signaling activates the PI3K/AKT cascade leading to inhibition of the transcription factor Forkhead box protein O1 (FOXO1) and reduced transcription of FOXO1 target genes (32). Ang2 generally functions as a TIE2 agonist in the context of cancer, presumably activating the PI3K/AKT cascade (32), a result we confirm here. Thus, cooperation between SHP099 and Ang1/2-TIE2 neutralization may derive from blocking non-overlapping pro-angiogenic cascades.

Endothelial cell death, vascular damage and involution of the tumor vasculature emerge as dominant effects of SHP099, associated with loss of vascular integrity and blood extravasation. These pathologies, uncommon consequences of anti-VEGF or anti Ang1/2 therapies, suggested participation of STAT signaling from SHP2 inactivation (10,45). To our knowledge, STAT activation is not a feature of Ang1/2 or VEGF neutralization. Thus, by activating STAT signaling, SHP099 may target VEGF- and Ang1/2-independent post-angiogenic tumor vessels.

The current results show that SHP2 inhibition represents a novel strategy to target endothelial cells in the tumor vasculature and raise the possibility that SHP2 inhibition could be particularly effective when applied more broadly to tumors where SHP2 functions as an oncogenic tyrosine phosphatase.

## Author contributions

GT conceived and directed the project; YW, OS and HO designed and executed the experiments; TA, JF, HK and MD helped with the execution of experiments; AT and MK contributed their expertise to image analyses and image quantification; DW performed bioinformatics analyses; and GT drafted the manuscript; all authors reviewed and contributed to text revisions.

## Acknowledgements

We thank Drs. I. Daar and D. Lowy for generously sharing reagents and helpful discussions; the animal facility personnel, particularly D. Gallardo; L. Lin, K. Stahl, K. Chen, C. Fedele, F. Luo, A. Sassano, C-P. Day, E. Perez-Guijarro, G. Merlino and the personnel of the Laboratory of Cellular Oncology for their help in various aspects of this project. Dr. Wang was a student at West China Medical School, Sichuan University during a period of this research. This work was supported by the intramural program of the Center for Cancer Research, NCI.

## Conflict of interest

The authors declare no conflict of interest

